# PEARL: Integrative multi-omics classification and omics feature discovery via deep graph learning

**DOI:** 10.1101/2025.05.19.654754

**Authors:** Quan Zhao, Jiawen Du, Muqing Zhou, Xu-Wen Wang, Quan Sun, Can Chen

## Abstract

Integrating multi-omics data provides valuable insights into biological processes by capturing information across multiple molecular layers, enabling a comprehensive understanding of complex diseases and driving advancements in precision medicine. However, existing computational methods for multi-omics integration face significant challenges, such as low reliability and poor generalizability, due to the high dimensionality and low sample size nature of omics data. To address these challenges, we present PEARL (Pearson-Enhanced spectrAl gRaph convoLutional networks), a novel deep graph learning method for biomedical classification and functional important omics features identification. PEARL leverages a simple yet effective learning architecture to achieve superior and robust performance in high-dimensional, low-sample-size multi-omics settings. Our results demonstrate that PEARL significantly outperforms existing state-of-the-art methods on both synthetic and real biomedical datasets. Furthermore, applied to Alzheimer’s disease (AD) brain multi-omics data, features prioritized by PEARL lead to functionally important genes that demonstrate significant enrichment in AD-related pathways. These findings highlight PEARL’s practical utility in biomedical research and its potential to enhance biological interpretability in multi-omics studies.

## Introduction

Advances in high-throughput sequencing technologies have revolutionized biomedical research by enabling the generation of diverse and extensive omics data, including genomic, transcriptomic, and epigenetic profiles, which are crucial for understanding complex diseases^1–5^. Integrating these multi-omics datasets provides a powerful approach to unraveling disease heterogeneity, uncovering functional important genes, and revealing dynamic molecular interactions underlying disease progression^6–10^. Moreover, as personalized medicine advances, multi-omics integration has become indispensable for refining disease diagnosis, tailoring individualized therapies, and discovering new therapeutic targets^11–13^. Therefore, developing robust multi-omics integration methods is salient to fully harness the potential of these diverse data types, leading to more accurate disease biology models, deeper understanding of molecular-level disease mechanisms, and more effective therapeutic strategies.

Numerous computational approaches have been developed for patient stratification, disease subtype classification, and drug target discovery through the integration of multi-omics data^7, 14, 15^. Among these, early integration methods typically combine features from multiple omics datasets into a single representation, while ensemble approaches aggregate predictions from separate classifiers, each trained individually on distinct omics data types^16–18^. However, these methods struggle to accommodate the intrinsic properties of omics data, such as low sample sizes and high dimensionality, and fail to capture the complex correlations across different types of omics data. The classical statistical method DIABLO^19^ employs canonical correlation analysis for multi-view integration but still faces difficulties in effectively modeling the nonlinear relationships across omics platforms with low sample sizes. With advanced computational capabilities, deep learning methods have emerged as powerful tools for multi-omics analysis. For instance, DeepKEGG^20^ introduces biologically informed hierarchical frameworks that leverage pathway-gene relationships for cancer recurrence prediction and functional genes discovery. DeepMo^21^ employs unsupervised learning approaches to stratify disease subtypes, while MOMA^22^ utilizes attention mechanisms to identify prognostic subtypes from multi-omics profiles. Despite their improved performance over traditional methods, these deep learning approaches often require substantial training data and computational resources, presenting barriers to clinical adoption.

Deep graph learning methodologies, such as graph convolutional networks (GCNs), have proven to be promising solutions for enhancing multi-omics integration. These methods excel at capturing both intra- and inter-omics relationships by representing samples as nodes in similarity networks. For example, MOGONET^23^ is a patient classification method that employs omics-specific GCNs constructed from sample similarity networks, integrating their predictions through a view correlation discovery network. Similarly, MOGDx^24^ utilizes a similarity network fusion to construct integrated patient networks, enabling robust disease classification even with heterogeneous data sources. MVGNN^25^ and MOGAT^26^ further advance these approaches by incorporating attention mechanisms to fuse multi-omics features and weigh the significance of molecular interactions. However, despite their potential, these methods often rely on highly complex architectures (e.g., graph attention networks), making them prone to overfitting and resulting in poor predictive performance when sample size is small.

To address these challenges, we propose PEARL (Pearson-Enhanced spectrAl gRaph convoLutional networks), a novel deep graph learning method for biomedical classification and functional important genes identification. PEARL leverages a simple but effective learning architecture, including weighted Pearson correlation, simple spectral graph convolutional networks, and a multi-layer perceptron, offering robust performance against sample size variations and noise commonly present in omics data. We systematically evaluate PEARL and other existing state-of-the-art multi-omics integration methods using both simulated and real biomedical datasets. Our results demonstrate that PEARL significantly outperforms existing methods, particularly under conditions of small sample sizes, while effectively identifying critical disease-associated functional genes, highlighting its potential for advancing precision medicine.

## Results

### Overview of PEARL

PEARL is a supervised multi-omics integration approach designed for biomedical classification and functional important omics features identification, effectively tackling the challenges of high dimensionality and limited sample sizes inherent in omics data. The framework consists of three key components: similarity network construction, feature refinement with GCNs, and feature integration (**Figure 1**).

**Fig. 1:**
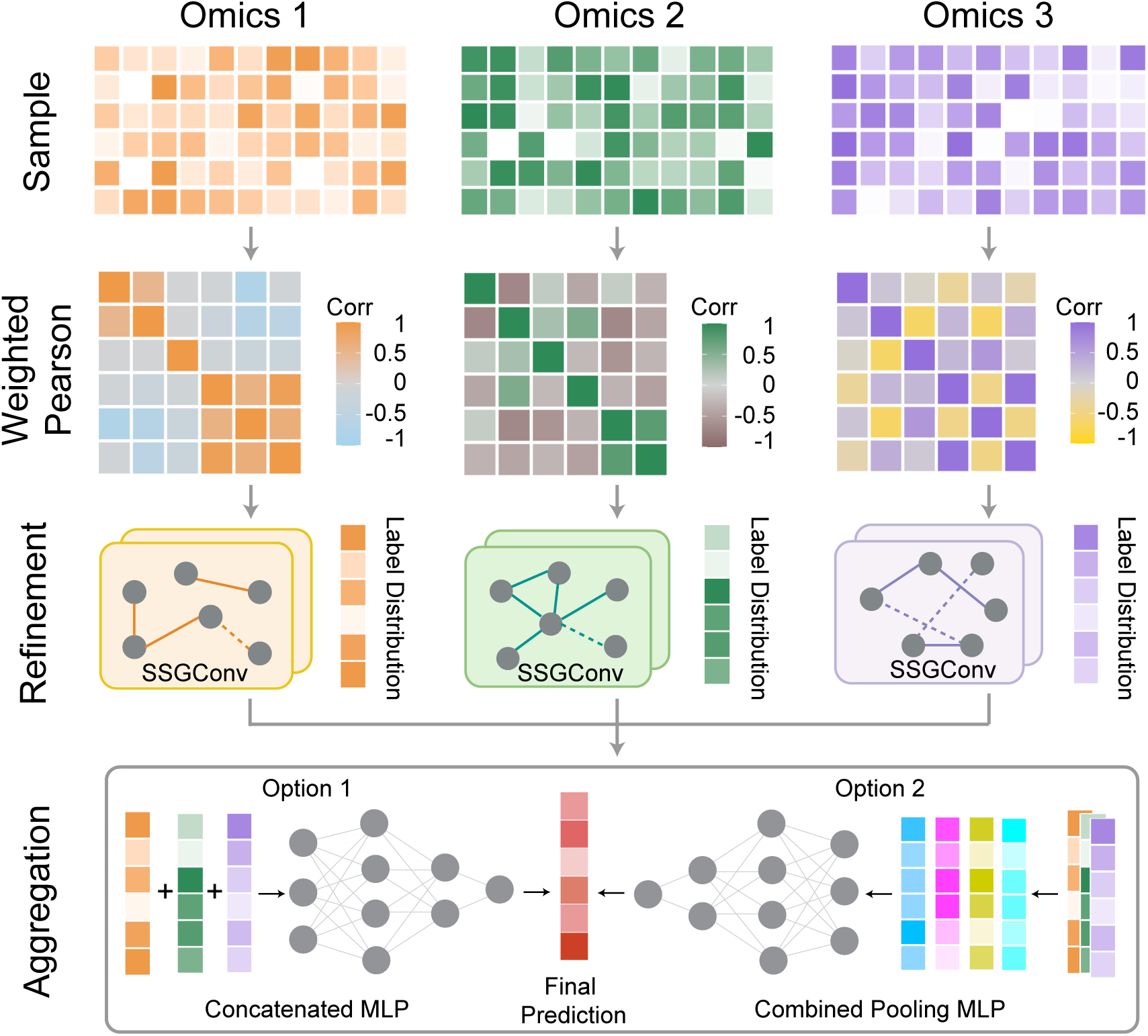
Overview of PEARL. PEARL is a novel multi-omics integration method based on deep graph learning for biomedical classification and functional important genes identification. Leveraging a simple yet effective architecture, PEARL first defines the graph structure for omics data using a weighted Pearson correlation similarity matrix. Omics-specific simple spectral graph convolutional networks are then employed for robust feature extraction from each view. For multi-view feature integration, PEARL offers two alternative strategies: a concatenated MLP or a combined pooling MLP, allowing for flexible fusion before the final prediction. The efficacy of PEARL has been validated across both simulated and real biomedical datasets.

First, PEARL constructs sample similarity networks for different omics data using weighted Pearson correlation^27^, where features with higher mean values have greater influence (see Methods). This weighting strategy is based on the assumption that features with lower mean values are more susceptible to noise, thus offering greater robustness than other network construction techniques, such as cosine similarity. Next, the similarity networks are represented as graphs for feature refinement using simple spectral graph convolutional networks (SSGConv)^28^. SSGConv is a simple yet effective graph learning architecture that enhances feature smoothing, reduces overfitting, and improves computational efficiency compared to traditional GCNs, making it particularly well-suited for high-dimensional, low-sample-size omics data^28^. The refinement process involves an initial SSGConv layer followed by ReLU activation and dropout for regularization, as well as a second SSGConv layer with activation. This dual-layer design allows the model to capture both local and global patterns in the data while minimizing overfitting. Finally, the refined features from each omics dataset are integrated using combined pooling and/or concatenation, and fed into a multilayer perceptron for the final prediction. The combined pooling method unifies various operations such as maximum, minimum, average, and max-min pooling, which collectively enhance feature representation by capturing extreme values, improving robustness to outliers, and boosting feature discrimination. Meanwhile, concatenation effectively integrates multiple feature sets by preserving all available information from diverse data sources. Detailed descriptions of each component can be found in the Methods section.

### PEARL demonstrates consistent superiority over existing methods across simulated scenarios

We first evaluated PEARL on synthetic data generated by InterSIM^29^, a method for generating synthetic multi-omics datasets that simulate real biological data. We considered three types of omics data, including methylation, gene expression, and protein data, and examined various cluster mean shifts and noise levels (i.e., difficulty levels) across sample sizes of 300, 500, and 800, resulting in a total of 9 synthetic datasets (Supplementary Note 2). PEARL was compared with state-of-the-art multi-omics integration methods MOGONET^23^, MOMA^22^, and MOGDx^24^ (Supplementary Note 1), as well as baseline classifiers, including K-nearest neighbor (KNN) and random forest (RF). We used accuracy, weighted average F1 score (F1 weighted), and macro-averaged F1 score (F1 macro) as evaluation metrics and reported the mean and standard deviation of those metrics from 30 random train-test splits with train-test ratio 8:2.

As illustrated in **Figure 2**, PEARL consistently outperforms all competing methods across varying sample sizes and difficulty levels, excelling in all evaluated metrics (accuracy, F1 weighted, and F1 macro). Notably, PEARL demonstrates exceptional robustness in low-sample-size settings (*n* = 300). For example, at the intermediate difficulty level (**Figure 2**b), PEARL achieves an average of 25% higher relative accuracy compared to the second-best method MOGONET (*P*-value = 4.69E-10, two-sided paired t-test). This performance advantage persists as sample sizes increase, with 30% and 32% higher accuracy compared to MOGONET in *n* = 500 and 800 scenarios (*P*-values = 4.18E-11 and 9.97E-14) under the same intermediate difficulty level. Additionally, PEARL effectively adapts to tasks of varying complexity, handling both noise-heavy scenarios with subtle patterns and simpler cases with clear class separations. Across all conditions, PEARL significantly surpasses alternatives like MOGONET, MOMA, and MOGDx with an average of 27%, 28%, and 27% higher accuracy. These results on synthetic data highlight the superiority of PEARL in various real-world data scenarios, especially given that multi-omics datasets are often limited by small sample sizes and high noise.

**Fig. 2:**
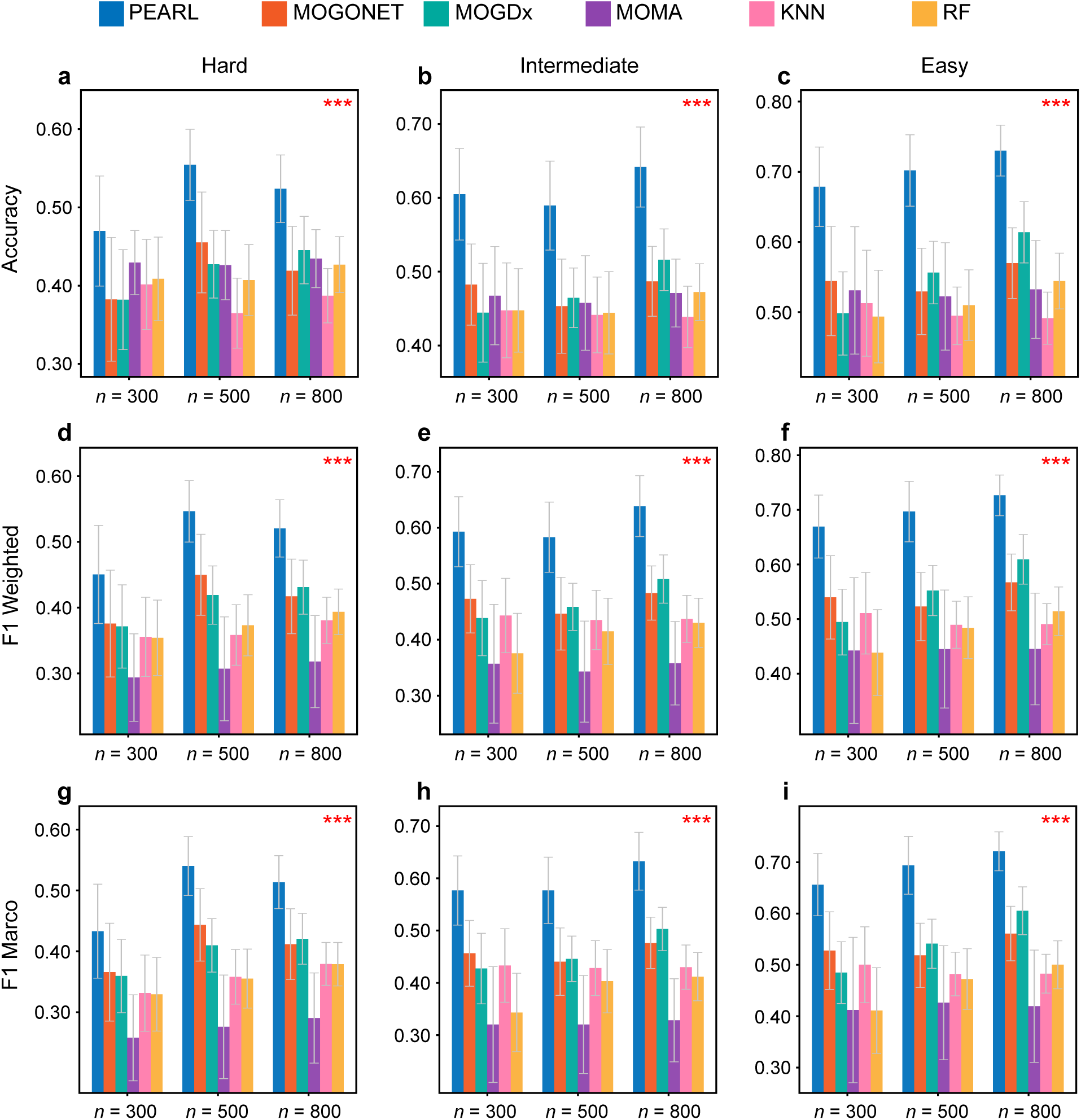
Performance comparison on synthetic datasets. Three sets of parameters were used with InterSIM to generate synthetic datasets. Hard refers to the cases when the cluster mean shift (delta) is equal to 0.1 and the noise level is equal to 0.4; Intermediate refers to the cases when the cluster mean shift is equal to 0.15 and the noise level is equal to 0.45; and then Easy refers to the cases when the cluster mean shift is equal to 0.2, and the noise level is equal to 0.5. For each case, we used three sample sizes (*n*) to generate synthetic datasets, which are 300, 500, and 800. We utilized Accuracy, F1 Weighted, and F1 Macro to measure the performance of PEARL, MOGONET, MOGDx, MOMA, KNN, and RF on the synthetic datasets. To assess statistical significance, we conducted paired t-tests comparing PEARL with the second-best method in each case. The resulting *P*-values are indicated with asterisks in the plots, where three asterisks denote a *P*-value less than 0.01 across all sample sizes.

### PEARL outperforms existing methods in various real datasets

We applied our method and other competing algorithms to two real datasets, including the Religious Orders Study/Memory and Aging Project (ROSMAP) for Alzheimer’s Disease (AD) patients vs. normal control (NC) classification^30, 31^, and BReast CAncer (BRCA)^32^ for breast invasive carcinoma PAM50 subtype classification. Both datasets contain three types of omics data: mRNA expression data (mRNA), DNA methylation data (meth), and miRNA expression data (miRNA). The numbers of features used for training are included in **Table 1**. Each dataset was randomly split into 80% training and 20% testing sets, repeated 30 times to ensure robust performance evaluation. For the binary ROSMAP dataset, we used accuracy (ACC), F1 score (F1), and area under the receiver operating characteristic curve (AUC) as evaluation metrics. Due to the multi-class characteristic of the BRCA data, we calculated ACC, average F1 score weighted by suppor (F1 weighted), and macro-averaged F1 score (F1 macro). We reported mean and standard deviation of each evaluation metrics across the 30 realizations.

**Table 1:** Summary of datasets. mRNA refers to mRNA expression data. meth refers to DNA methylation data. miRNA refers to miRNA expression data. The ROSMAP dataset is for the classification of Alzheimer’s disease (AD) patients vs. normal control (NC). The BRCA dataset is for breast invasive carcinoma (BRCA) PAM50 subtype classification with normal-like, basal-like, human epidermal growth factor receptor 2 (HER2)-enriched, Luminal A, and Luminal B subtypes.

| **Dataset** | **Categories** | **Number of features**<br>mRNA, meth, miRNA | **Number of features**<br>training mRNA, meth,<br>miRNA |
| --- | --- | --- | --- |
| **ROSMAP** | NC: 169, AD: 182 | 55,889, 23,788, 309 | 200, 200, 200 |
| **BRCA** | Normal-like: 115,<br>Basal-like: 131,<br>HER2-enriched: 46,<br>Luminal A: 436,<br>Luminal B: 147 | 20,531, 20,106, 503 | 1000, 1000, 503 |

Our results show that PEARL consistently outperforms all other methods in both BRCA and ROSMAP datasets (**Figure 3**), with notable improvements in all metrics. In the BRCA dataset with a challenging situation of classifying five different subtypes of breast cancer (**Figure 3**a-c), PEARL outperforms MOGONET by more than 2% in Accuracy, more than 5% in F1 weighted, and more than 8% in F1 macro (*P*-value = 3.16E-04, 3.49E-07, 5.32E-07, respectively), while maintaining smaller standard deviations (0.0432 for PEARL compared to 0.0610 for MOGONET in F1 macro). The reduced standard deviations in PEARL’s results also suggest more consistent and reliable predictions, a critical factor for clinical implementation where reproducibility is essential. In the ROSMAP dataset (**Figure 3**d-f), which presents the challenging task of distinguishing AD patients from normal controls, PEARL demonstrates remarkable improvements compared to its closest competitor MOGONET, achieving gains of more than 2% in Accuracy and AUROC, more than 3% in F1 score (*P*-value = 2.35E-03, 1.22E-03, 5.09E-05, respectively). These improvements are particularly meaningful in the context of AD diagnosis, where even small increases in accuracy can potentially translate to earlier and more reliable disease detection, potentially enabling earlier intervention strategies.

**Fig. 3:**
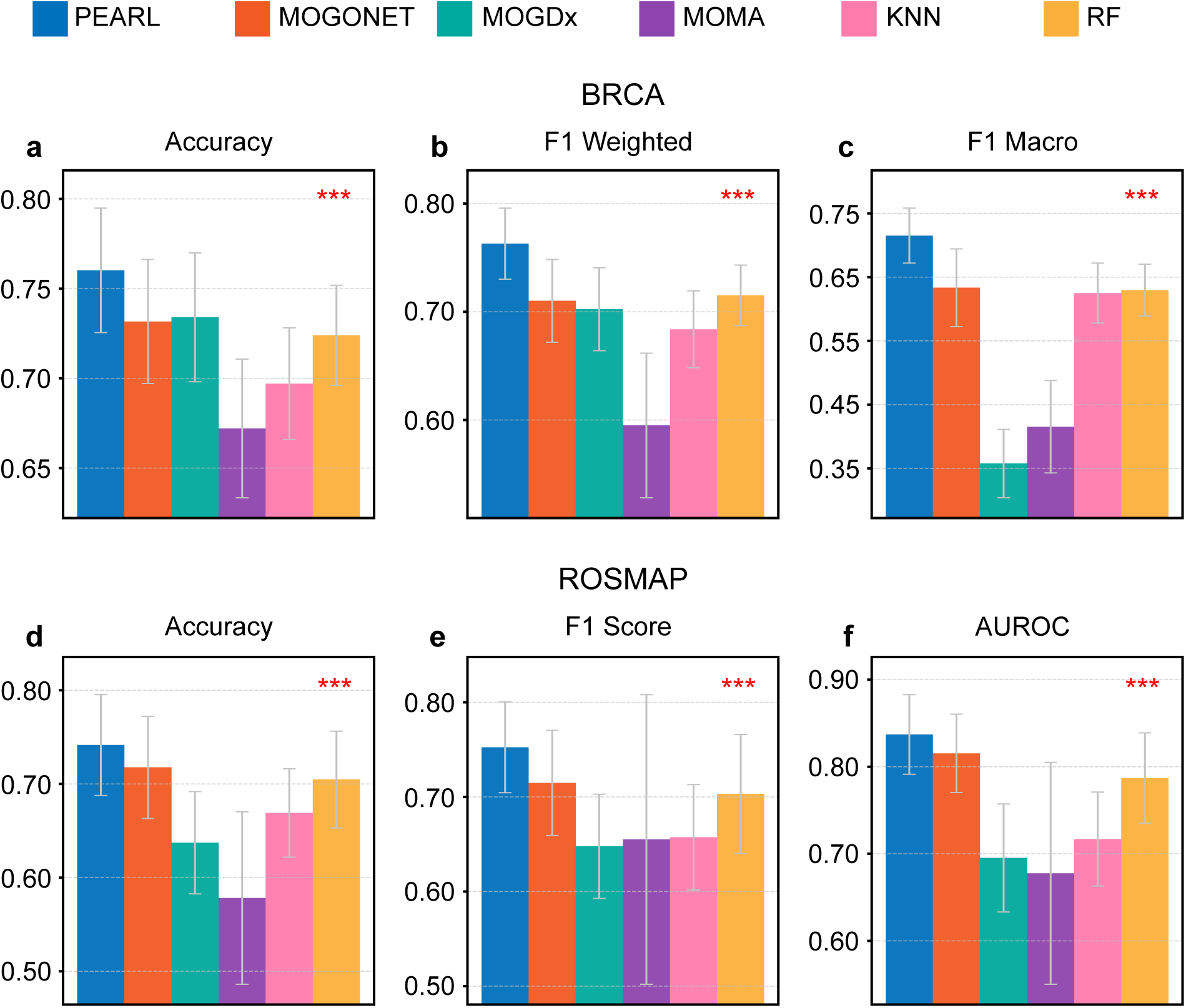
Performance comparison on real datasets. We compared the performance of PEARL with four other methods, MOGONET, MOGDx, MOMA, KNN, and RF, on the BRCA dataset and the ROSMAP dataset. We used Accuracy, F1 Weighted, and F1 Macro as metrics on the BRCA multi-view dataset. For the ROSMAP binary dataset, we calculated Accuracy, F1 Score, and AUROC as evaluation metrics. To assess statistical significance, we conducted paired t-tests comparing PEARL with the second-best method in each case. The resulting *P*-values are indicated with asterisks in the plots, where three asterisks denote a *P*-value less than 0.01.

All methods were run with their default parameter configurations to ensure consistency in comparative evaluation in above analysis. To evaluate the impact of parameter tuning on existing methods, we adopted a grid search strategy, and the results confirm the superiority of PEARL (Supplementary Figure 1). We further performed an ablation study to examine the impact of our key innovations in PEARL (Supplementary Note 3). The results demonstrate that removing any one of our key components, including weighted Pearson correlation, SSGConv, or feature integration, causes a stepwise decline in all performance metrics, confirming that each innovation is essential for PEARL’s best results (Supplementary Figure 2).

### Identification of Disease-Associated Genes and Pathways

Finally, we investigated the key omics features that contributed the most to the prediction (**Tables 2 and 3**). Details of the identification process are provided in the Methods section. In the ROSMAP dataset, we chose the top 50 features with highest importance scores, across mRNA, DNA methylation and miRNA data. We annotated the cpg sites to their nearest gene using Illumina MethylationEPIC v2.0 manifest file. For miRNA, they were filtered to those present in the TargetScan human database^33^ and further mapped to corresponding miRNA families. Further, the target genes were extracted from the TargetScan’s target predictions. All gene symbols were subsequently converted to ENSEMBL gene IDs using the *org.Hs.eg.db*^34^ annotation database. Overall, these 50 features correspond to 2,861 genes. Then, we performed enrichment analysis for the functional genes identified in ROSMAP, using both Gene Ontology (GO) and Kyoto Encyclopedia of Genes and Genomes (KEGG) databases^35^. GO terms within three ontology categories are considered: biological process (BP), cellular components (CC), and molecular function (MF). To account for multiple testing, false discovery rate (FDR) correction was applied and the adjusted *P*-values were reported. We found these genes were enriched for several GO terms that have been reported in other studies to be associated with Alzheimer’s disease. Notable examples include small GTPase mediated signal transduction (biological process, GO:0007264, adjusted *P*-value = 1.07E-16, **Figure 4**)^36^, axongenesis (biological process, GO:0007409, adjusted *P*-value = 5.11E-13, **Figure 4**)^37^, axon development (biological process, GO:0061564, adjusted *P*-value = 2.56E-12, **Figure 4**)^38^, glutamatergic synapse (cellular composition, GO:0098978, adjusted *P*-value = 1.36E-23, Supplementary Figure 3)^39^, and protein serine/threonine kinase activity (molecular function, GO:0004674, adjusted *P*-value = 2.98E-9, Supplementary Figure 4)^40^. The full GO enrichment results are provided in Supplementary Tables 3-5.

**Fig. 4:**
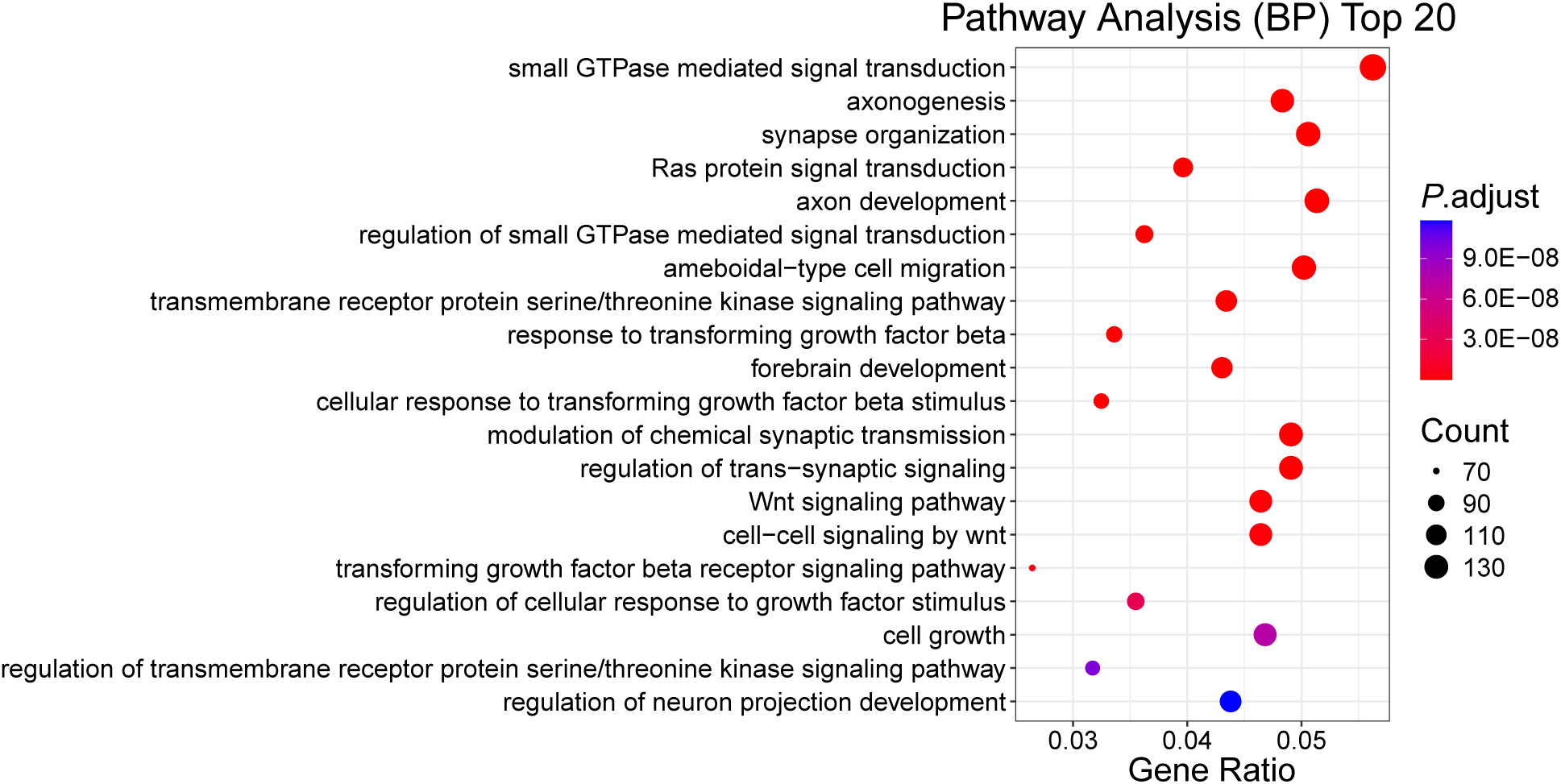
Top 20 enriched biological processes from pathway analysis ordered by adjusted *P* -value. Each dot represents a biological process, with the x-axis showing the gene ratio (the ratio of genes associated with a given term to the total number of genes analyzed), and the y-axis listing the description of each biological process. Color corresponds to the adjusted *P*-value, with more intense red indicating higher significance. Dot size reflects the gene count. Several notable significantly enriched pathways include processes related to GTPase-mediated signaling, axongenesis and axon development.

**Table 2:** Feature identification results from ROSMAP by PEARL.

| **Omics data type** | **Features** |
| --- | --- |
| **mRNA**<br>**expression** | ENSG00000117266.11, ENSG00000155980.6,<br>ENSG00000105643.3, ENSG00000204219.5,<br>ENSG00000182310.7, ENSG00000168743.8,<br>ENSG00000071054.10, ENSG00000102032.8,<br>ENSG00000166682.6 |
| **DNA methylation** | cg07232688, cg22518733, cg27091787, cg25736482,<br>cg14387505, cg12447832, cg08418111, cg02008416,<br>cg06946880, cg10157098, cg24765079, cg04983977, cg14847483 |
| **miRNA**<br>**expression** | hsa-miR-517c_hsa-miR-519a, hsa-miR-424, hsa-miR-216a, hsa-miR-651, hsa-miR-127-3p, hsa-miR-640, hsa-miR-199b-5p, hsa-miR-199a-5p |

**Table 3:** Feature identification results from BRCA by PEARL.

| **Omics data type** | **Feature** |
| --- | --- |
| **mRNA expression** | SOX11—6664, HPDL—84842, CNGA1—1259, KIF2C—11004, AR—367, CAPN13—92291, CDC6—990, KIF14—9928, PTCHD1—139411, C1orf64—149563, PHYHD1—254295 |
| **DNA methylation** | C14orf72, OR1J4, TM4SF18, LEMD1, ROPN1, ATP8B1, C21orf82, GPR37L1, KDM2B, ATP10B, LGALS2 |
| **miRNA expression** | hsa-mir-187, hsa-mir-9-1, hsa-mir-9-2, hsa-mir-203, hsa-mir-21, hsa-mir-184, hsa-mir-944, hsa-mir-551b |

The KEGG pathway enrichment analysis also identified several important pathways that are known to be critically related to AD (Supplementary Table 6). The two most notable pathways are axon guidance (adjusted *P*-value = 2.06E-13, Supplementary Figure 5) and MAPK signaling pathway (adjusted *P*-value = 1.40E-10, Supplementary Figure 5). Axon-guidance molecules are playing import roles in occurrence and development of AD by influencing in connectivity and repair mechanism^41^, and the activate MAPK signaling pathway has been considered to contribute to AD via multiple mechanism like neuronal apoptosis and transcriptional and enzymatic activation of *β*- and *γ*-secretases^42^. These findings demonstrate that PEARL effectively identifies trait-associated functional genes from multi-omics data, supporting its utility in uncovering biologically meaningful insights.

## Discussion

Multi-omics integration plays crucial roles in understanding disease mechanisms, providing pivotal perspectives in disease diagnosis and prognosis. In this study, we present PEARL, a novel framework for multi-omics data integration and analysis that advances the field through several key innovations. Our comprehensive evaluation demonstrates PEARL’s superior performance across both synthetic and real-world datasets, while also providing biologically meaningful insights through its feature identification capabilities.

PEARL’s success can be attributed to several major technical innovations. First, the weighted Pearson correlation approach in the pre-processing stage represents a significant advancement over traditional similarity measures. By incorporating a position-based weighting scheme, PEARL captures the varying importance of different molecular features, reflecting the biological reality that not all features contribute equally to disease mechanisms. This builds upon insights from previous studies showing that representing data as patient similarity networks can retain information and have better performance compared to standard Euclidean methods^43, 44^. Second, the use of simple but effective SSGConv layers in PEARL’s GCN architecture provides another crucial advantage. Unlike traditional approaches that rely on simple feature concatenation^45^ or majority voting for network integration^46^, SSGConv’s ability to balance between feature preservation and neighborhood aggregation is especially valuable in handling the noise and heterogeneity inherent in multi-omics data. This is reflected in the consistently smaller standard deviations observed in our results, particularly in the ROSMAP dataset. Third, the dual integration strategies, combined pooling and concatenation MLP, demonstrate PEARL’s robustness to different classification scenarios. This flexibility addresses limitations identified in existing methods^47^ where integrative analysis often struggles with different types of classification tasks. Each component of PEARL contributes to its superiority over competing methods, validated through our results with enhanced performance for both synthetic and real datasets.

PEARL addresses several limitations of existing approaches. Unlike MOGONET, which relies on a fixed integration framework, PEARL offers more flexible handling of different data types and classification scenarios. Compared to MOMA, which faces challenges with complex optimization in multi-task learning settings that can create negative transfer effects, PEARL’s architecture avoids these conflicts while maintaining computational efficiency. Similarly, while MOGDx effectively handles missing data through network fusion, it lacks the sophisticated feature weighting mechanisms present in PEARL. The weighted Pearson correlation and SSGConv architecture in PEARL provide a more robust framework for high-dimensional, low-sample-size data compared to these methods. Furthermore, PEARL’s adaptive integration approach delivers superior performance across varying classification scenarios without extensive parameter tuning, offering better scalability and robustness as demonstrated in our synthetic data experiments across different sample sizes.

PEARL not only achieves higher accuracy in classification tasks, but also has the ability to identify biologically meaningful important features from various omics data, providing valuable insights into disease mechanisms. Like recent successful methods^48^, PEARL effectively integrates multiple omics data types while maintaining interpretability. In the ROSMAP data analysis, the underlying genes of prioritized omics features by PEARL show significant enrichment in AD-related pathways, particularly axon guidance and MAPK signaling, consistent with current understanding of AD pathophysiology. Of particular note is PEARL’s ability to identify functionally important pathway associations. Like recent network-based approaches^49^, PEARL excels at capturing strong latent relationships between samples. However, PEARL goes beyond simple network fusion by incorporating weighted correlations and advanced graph convolution techniques, allowing for more nuanced feature importance estimation.

While PEARL demonstrates significant advantages, several areas warrant further investigation. First, the current implementation focuses on classification tasks; extending the framework to handle regression problems could broaden its clinical applications. This could potentially build upon approaches used in previous studies for survival prediction^50^. Second, the computational complexity of PEARL, while manageable for current data sizes, may present challenges with larger-scale multi-omics studies. This reflects a common problem in the field^51^, and future work could explore optimization strategies or distributed computing approaches to improve scalability. Last, another promising direction would be to integrate PEARL with emerging approaches in multi-omics integration, such as canonical correlation analysis methods^52^ or advanced group-regularized techniques^53^, which may provide additional insights into cellular heterogeneity and tissue-specific patterns in disease progression.

In summary, PEARL represents a significant advancement in multi-omics data integration and analysis. Its superior performance across various datasets, coupled with its ability to identify biologically meaningful features, positions it as a valuable tool for precision medicine applications. Building on insights from previous unsupervised integration methods^54^, PEARL extends these capabilities to supervised learning while maintaining robust performance in the presence of noise. As multi-omics studies continue to grow in scale and complexity, we anticipate PEARL will become increasingly valuable for understanding disease mechanisms and advancing personalized medicine approaches.

## Methods

### Network construction

PEARL employs weighted Pearson correlation^27^ to assign feature-specific weights and compute a correlation network for each view. Unlike the traditional Pearson correlation coefficient, the weighted version incorporates a tailored weighting scheme for each feature, effectively addressing a key challenge in multi-omics data analysis, namely, that not all features contribute equally to underlying biological processes. Our approach leverages feature means as biologically-informed weights, based on the observation that features with higher average expression or activity levels often reflect greater biological relevance. Specifically, given two samples with feature vectors **x** and **y**, and a feature weight vector **w** derived from the mean value of each feature across all samples, we first compute the weighted feature representations as **x**_weighted_ = **w** ⊙ **x** and **y**_weighted_ = **w** ⊙ **y**, respectively, where ⊙ denotes element-wise multiplication. We then center the weighted features by subtracting their respective means, i.e., **x**_centered_ = **x**_weighted_ − mean(**x**_weighted_) and **y**_centered_ = **y**_weighted_ − mean(**y**_weighted_). Finally, the weighted Pearson correlation between the two samples is computed as

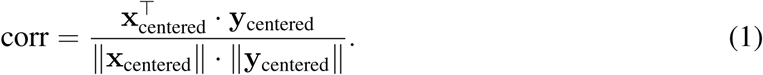

Using this simple yet biologically meaningful formulation, we effectively construct a sample similarity network for each modality that captures latent relationships between samples.

### Feature refinement

PEARL utilizes SSGConv (simple spectral graph convolutional network) layers^28^ to refine sample features over the constructed similarity networks. SSGConv is a simple but sophisticated spectral-based deep learning architecture that is particularly well-suited for multi-omics data analysis scenarios characterized by high dimensionality and limited sample sizes. For each omics-specific similarity network, the SSGConv layer is defined as

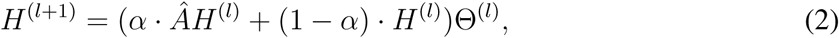

where *Â* is the normalized adjacency matrix of the network with added self-loops, *H*^(^*^l^*^)^ is the feature matrix at layer *l* (*H*^(0)^ represents the original omics data), Θ^(^*^l^*^)^ is the learned parameter matrix at layer *l*, and *α* is a weighting factor that balances the smoothness of the neighborhood aggregation with the retention of the previous features. SSGConv can be implemented efficiently using sparse matrix operations, crucial for handling high-dimensional omics data. In PEARL, we employ a two-layer SSGConv architecture to refine each omics feature representation over its corresponding similarity network, incorporating ReLU as the nonlinear activation function and dropout for regularization. The use of SSGConv equips PEARL with a robust framework for handling high-dimensional, low-sample-size multi-omics data, enabling effective feature learning while ensuring computational efficiency.

### Feature integration

PEARL offers two feature integration strategies: combined pooling and concatenation. The combined pooling approach integrates features from multiple omics views using a variety of pooling operations, such as maximum, minimum, average, and difference pooling, followed by a fully connected multi-layer perceptron (MLP) for classification. Given the refined features from *M* views {*h*^(1)^*, h*^(2)^*, … , h*^(^*^M^*^)^} for each sample, the pooling operations can be mathematically defined as

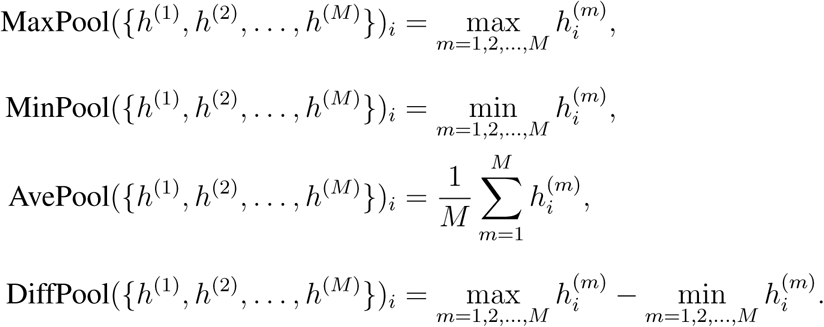

The pooled features are concatenated to form a unified feature vector for each sample, denoted by *h*. This unified feature vector is then passed through an MLP with a final output dimension of two for binary classification, i.e.,

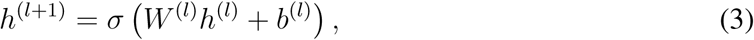

where *W* ^(^*^l^*^)^ and *b*^(^*^l^*^)^ represent the weights and biases of the *l*th layer, respectively, and *σ* denotes the activation function (e.g., Softmax). The second pooling approach, i.e., concatenation, is relatively simpler, where it directly concatenates the refined features from different omics views and feeds them into an MLP for final classification. The two pooling strategies in PEARL provide a comprehensive yet flexible framework for integrating features across diverse omics types.

### Feature identification

Beyond its prediction power, PEARL also has the ability to identify the important features of omics data that contribute the most to prediction. Specifically, for a given feature, we first set its value vector to zero to simulate the absence of this feature as a perturbed dataset. This dataset is then processed as usual and is used to train a perturbed prediction model. We define the importance score of this feature by calculating the difference in model performance metrics (F1 score or F1 macro) between the perturbed model and the baseline model that includes this feature. We note that each omics data may contribute to the prediction unequally. Thus, we normalize the importance scores across all the features within each omics layer. In our experiments, we performed five repeats of each dataset and calculated the average of feature importance scores to account for randomness of training process.

### GO and KEGG Enrichment Analysis

To explore the biological functions and pathways associated with the functional important genes identified by PEARL, we first used MethylationEPIC v2.0 annotation from Illumina and *TargetScan* to map the corresponding genes from epigeneticmarkers and miRNA. After obtaining the gene set, GO and KEGG pathway enrichment analysis were conducted with the *enrichGO* and *enrichKEGG* functions, respectively, from the *clusterProfiler* R package^55^. To account for multiple testing, FDR was calculated by adjusting *P*-values using the Benjamini-Hochberg method. GO terms and pathways with adjusted *P*-values *<* 0.05 are considered to be significant.

## Data Availability

The ROSMAP dataset was obtained from AMP-AD Knowledge Portal (https://adknowledgeportal.synapse.org/). Omics data of BRCA was obtained from The Cancer Genome Atlas Program (TCGA) through Broad GDAC Firehose (https://gdac.broadinstitute.org/). PAM50 breast cancer subtypes of TCGA BRCA patients were obtained through the TCGAbiolinks R package (v2.12.6, http://bioconductor.org/packages/release/bioc/html/TCGAbiolinks.html). Source data are provided with this paper.

## Code Availability

The source code of our computational framework is available at https://github.com/zqq121017/PEARL.

## Competing Interests Statement

The authors declare that they have no competing interests.

## Notes

### Competing Interest Statement

The authors have declared no competing interest.

