## Supplementary Material for "PEARL: Integrative multi-omics classification and omics feature discovery via deep graph learning"

Quan Zhao *et al.*

### Supplementary Note 1. Methods Compared with PEARL

##### MOGONET (Multi-Omics Graph cOnvolutional NETworks)

MOGONET[1] is a supervised integration framework designed to enhance biomedical classification and biomarker discovery by integrating multi-omics data. Starting with preprocessing to eliminate noise and redundant features, it employs Graph Convolutional Networks (GCNs) to develop detailed omics-specific representations and uncover significant cross-omics correlations. This preparatory stage ensures that each dataset is optimally prepared for more effective and accurate analysis. Each type of omics data is processed through its respective GCN, utilizing a weighted sample similarity network constructed via cosine similarity. This network is defined by the equation:

$$A_{ij} = \begin{cases} \frac{\mathbf{x}_i \cdot \mathbf{x}_j}{\|\mathbf{x}_i\| \|\mathbf{x}_j\|} & \text{if } i \neq j \text{ and } \frac{\mathbf{x}_i \cdot \mathbf{x}_j}{\|\mathbf{x}_i\| \|\mathbf{x}_j\|} \geq \epsilon \\ 0 & \text{otherwise} \end{cases} \tag{1}$$

where $x_i$ and $x_j$ are the feature vectors of nodes i and j, respectively. This formula refines the adjacency matrix by only considering node pairs with a cosine similarity above a threshold $\epsilon$, ensuring that only significant relationships are modeled.

The transformation defining each layer of the GCN is given by $H^{(l+1)} = \sigma\left(AH^{(l)}W^{(l)}\right)$, which extracts complex features from the data. To optimize the learning phase, the adjacency matrix $A$ is modified to $\tilde{A} = D^{-\frac{1}{2}}(A + I)D^{-\frac{1}{2}}$, ensuring efficient network operations and effective feature integration. The framework further enhances data interpretation through a cross-omics discovery tensor $C_j$ in $\mathbb{R}^{c \times c \times c}$, facilitating comprehensive analysis of label correlations across different omics layers. This tensor is then reshaped and processed by the View Correlation Discovery Network (VCDN) for final predictions.

PEARL builds upon and extends the foundation established by MOGONET and introduces several key innovations that enhance both the theoretical framework and practical performance.

The fundamental architectural elements shared between PEARL and MOGONET[1] provide a strong foundation for multi-omics analysis. Both frameworks implement a multi-component pipeline that processes omics data through preprocessing, omics-specific learning, and integration stages. They utilize graph-based approaches for capturing sample relationships within each omics type and provide end-to-end trainable frameworks that jointly optimize omics-specific and integration components. Additionally, both systems are designed to handle multiple types of omics data, including mRNA expression, DNA methylation, and miRNA expression.

PEARL advances beyond MOGONET[1] through several significant enhancements to this architecture. In the preprocessing stage, PEARL introduces a sophisticated weighted Pearson correlation-based approach with a novel position-based weighting scheme. This innovation provides more nuanced feature importance estimation that better accounts for the varying biological significance of different molecular markers. The preprocessing pipeline also incorporates improved mechanisms for handling noise and redundancy in the input data. A major architectural advancement in PEARL is the replacement of traditional graph convolution with Simple Spectral Graph Convolution (SSGConv). This modification achieves a better balance between feature preservation and neighborhood aggregation, enhancing the framework's ability to capture both local and global patterns in molecular data. The SSGConv architecture provides more flexible spectral learning compared to MOGONET's fixed spectral bands approach. PEARL also introduces more sophisticated integration strategies compared to MOGONET's universal VCDN-based integration. The framework implements two distinct integration approaches: Combined Pooling MLP for multi-class problems and Enhanced Concatenation MLP for binary classification. This adaptive approach allows for more flexible handling of different classification scenarios, particularly beneficial in complex disease classification tasks. The integration modules are specifically designed to improve cross-omics correlation learning while maintaining computational efficiency.

The technical implementation differences between PEARL and MOGONET[1] extend to network architecture and feature processing. While MOGONET[1] employs a standard cosine similarity-based approach for feature processing, PEARL utilizes weighted correlation with position-based importance, providing more nuanced relationship metrics between samples. The network architecture in PEARL leverages SSGConv with adaptive spectral learning, contrasting with MOGONET's traditional graph convolution approach. PEARL's enhanced framework also includes comprehensive regularization techniques, including residual connections and sophisticated batch normalization strategies. These improvements work in concert with robust dropout mechanisms to prevent overfitting while maintaining model stability. Together, these technical advancements create a more robust and adaptable framework while preserving the interpretability that made MOGONET[1] successful.

**MOGDx (Multi-Omic Graph Diagnosis)**

MOGDx[2] is a sophisticated tool designed to integrate multi-omic data for the classification of heterogeneous diseases. It utilizes a network taxonomy that combines patient similarity networks through the Similarity Network Fusion (SNF) algorithm, enriches these networks with reduced vector representations of genomic data, and employs a Graph Convolutional Network (GCN) for classification tasks. MOGDx[2] excels in integrating multiple omic datasets—including genomic, transcriptomic, and proteomic data—to deliver state-of-the-art classification results for diverse diseases. This network-based approach efficiently manages and integrates data, adeptly handling missing data through network fusion, and demonstrates superior performance in identifying relevant multi-omic markers and achieving high classification accuracy across various datasets.

A patient similarity matrix is created for each modality. The Pearson correlation coefficient between the extracted features is used as a measure of similarity where suitable, otherwise Euclidean distance is used. The equations are given by:

$$r = \frac{\sum (x_i - \bar{x})(y_i - \bar{y})}{\sqrt{\sum (x_i - \bar{x})^2 \sum (y_i - \bar{y})^2}} \tag{2}$$

$$r(p, q) = \sqrt{\sum_{i=1}^{n} (q_i - p_i)^2} \tag{3}$$

The K-Nearest Neighbours (KNN) algorithm is used to build the graph with edges created between the 15 nearest neighbors. SNF is applied to fuse the graphs into a single network representing the full spectrum of the underlying data. SNF allows complementary information to be shared between modalities and is effective in identifying novel relationships between patients. It also integrates missing patient samples inherently by complementing a missing edge in one modality with the same relationship from others.

The core computational method in MOGDx[2] involves the use of a GCN, where each layer's transformation is defined by the equation:

$$H^{(l+1)} = \sigma\left(\tilde{A} H^{(l)} W^{(l)}\right) \tag{4}$$

where $H^{(l)}$ represents the matrix of node features at layer $l$, $\tilde{A}$ is the adjacency matrix modified to include self-connections, $W^{(l)}$ is the weight matrix for that layer, and $\sigma$ is the activation function.

**MOMA (Multi-task Attention Learning Algorithm for Multi-omics Data )**

MOMA[4] is a supervised learning method designed to integrate multi-omics data for biomedical classification and biological interpretation. It employs a geometric approach where genes and modules are vectorized, and uses an attention mechanism to focus on related modules across different omics datasets. The framework consists of three main stages: module encoding, module attention, and multi-task learning.

In the module encoder stage, MOMA utilizes a fully connected layer that connects features of each omics data to modules. Each module is represented as a vector, where the weights of the fully connected layer $\theta^j_{module}$ module represent associations between features and modules of the $j$th omics data. Given a training sample $\{x^j, y\}$, where $x^j$ denotes the sample under the $j$th omics of $J$ omics datasets and $y$ is the corresponding label, the module vectors $M^j$ are defined as:

$$M^j(x^j) = f^j_{module}(x^j; \theta^j_{module}) \in \mathbb{R}^{N^j \times D}, \tag{5}$$

where $\theta_module$ denotes the weights of $f_{module}$, $N^j$ is the number of modules of $j$th omics data, and $D$ is the dimension of the module vector.

The module attention mechanism focuses on modules with high similarity between each omics data module using cosine similarity. Let $Att$ denote the module attention matrix between the module vectors of two omics datasets. The attention matrix element $Att_{lk}$ is computed as:

$$Att_{lk}(M^i, M^j) = \frac{\exp(\cos(M_l^i, M_k^j))}{\sum_{k=1}^{N_j} \exp(\cos(M_l^i, M_k^j))} \tag{6}$$

where $M_l^i$ and $M_k^j$ are the $l$th module vector of $i$th omics data and the $k$th module vector of $j$th omics data, respectively. The updated module vector is:

$$\mathrm{Att}_M^j(x^{j'}) \;=\; \left[\left(\mathrm{Att}(M^j, \overline{M}^{j'})\right)^T M^j\right], \quad \text{s.t.} \quad \overline{M}^{j'} \in \{\, M \mid M \neq M^{j'} \} \tag{7}$$

Finally, fully connected layers are applied to yield the final probabilities for each label. The loss L is set to the cross-entropy error between the gold label and task-specific outputs:

$$L = -\sum_{j=1}^{J}\sum_{c=1}^{C} y_c \cdot \log(f_{fc}^j(Att^{M_j}(x^j); \theta_{fc}^j)) + \lambda \sum W^2, \text{ s.t. } W \in \{\theta_{module}, \theta_{fc}\}, \tag{8}$$

where $C$ represents the total number of classes, an L2-norm penalty with regularization parameter $\lambda$ is used to avoid overfitting, and $f_{fc}^j$ consists of multiple fully connected layers for jth omics data.

**KNN (K-Nearest Neighbor)**

The k-Nearest Neighbors algorithm represents a non-parametric, instance-based learning approach. For a given test sample x, KNN identifies the k training samples closest to x in the feature space and makes predictions based on majority voting among these neighbors. The proximity between samples is calculated using Euclidean distance in the feature space:

$$d(x, x_i) = \sqrt{\sum_{j=1}^{n} (x_j - X_{ij})^2} \tag{9}$$

where $x$ and $x_i$ are two samples, $n$ is the number of features, and $x_j$ and $X_{ij}$represent their $j$-th features respectively. The predicted class $\hat{y}$ for sample x is determined by:

$$\hat{y} = argmax_{c \in C} \sum_{i \in N_k(x)} I(y_i = c) \tag{10}$$

where $C$ is the set of possible classes, $N_k(x)$ represents the $k$ nearest neighbors of $x$, $y_i$ is the class label of the $i$-th neighbor, and $I()$ is the indicator function.

##### RF (Random Forest Classifier)

Random Forest utilizes an ensemble of decision trees, where each tree is trained on a bootstrap sample of the training data. At each node in a decision tree, the optimal split is determined using Gini impurity rather than the previously stated entropy-based information gain. The Gini impurity for a set of samples S is calculated as:

$$G(S) = \sum_{i=1}^{c} p_i(1 - p_i) \tag{11}$$

where $p_i$ is the proportion of samples in class $i$, and $c$ is the number of classes. For each split, the feature and threshold that maximize the reduction in Gini impurity are selected:

$$\Delta G = G(S) - \frac{|S_L|}{|S|}G(S_L) - \frac{|S_R|}{|S|}G(S_R) \tag{12}$$

where $S_L$ and $S_R$ are the left and right child nodes after the split, respectively. The final prediction is made through majority voting across all trees:

$$\hat{y} = argmax_{c \in C} \sum_{t=1}^{T} I(h_t(x) = c) \tag{13}$$

where T is the number of trees and $h_t(x)$ is the prediction of the $t$-th tree for sample $x$.

#### Supplementary Note 2. Synthetic datasets

##### Generation of synthetic datasets with InterSIM

InterSIM[3] is a specialized R package designed to simulate multiple interrelated genomic datasets that maintain realistic relationships both within and between different omics data types. Originally developed to aid in the evaluation of integrative clustering methods, InterSIM[3] generates synthetic data based on real molecular profiles from The Cancer Genome Atlas (TCGA) ovarian cancer studies. The package's key strength lies in its ability to preserve important biological relationships, such as the correlation between CpG methylation and downstream gene expression, and between gene expression and corresponding protein levels. InterSIM also maintains the complex covariance structures within each omics layer that are typically observed in real biological data.

For our synthetic data evaluation, we utilized InterSIM[3] to generate a comprehensive set of synthetic datasets. InterSIM is a specialized R package designed to simulate multiple interrelated genomic datasets that maintain realistic relationships both within and between different omics data types. The package generates synthetic data based on real molecular profiles from The Cancer Genome Atlas (TCGA) ovarian cancer studies, preserving important biological relationships such as the correlation between CpG methylation and downstream gene expression, and between gene expression and corresponding protein levels. InterSIM also maintains the complex covariance structures within each omics layer that are typically observed in real biological data.

We generated nine synthetic datasets by combining three different sample sizes (300, 500, and 800) with three different configurations of cluster mean shifts (delta values) and noise levels. For each dataset, samples were distributed across three clusters in proportions of 30%, 30%, and 40%. The configurations were structured to represent varying levels of classification difficulty: hard cases with delta = 0.1 and noise level = 0.4, intermediate cases with delta = 0.15 and noise level = 0.45, and easy cases with delta = 0.2 and noise level = 0.5. Each dataset included three types of omics data: DNA methylation, gene expression, and protein expression. The simulation process involved multiple coordinated steps. Initial data generation utilized InterSIM with independent covariance structures for each omics type and a proportion of 0.2 for differentially methylated positions. The package automatically handled the generation of corresponding differentially expressed genes and proteins based on the established biological relationships from TCGA data. To simulate real-world variability, we added Gaussian noise to each omics layer, with the noise level scaled relative to the standard deviation of each dataset. This step helped simulate the technical and biological variation commonly observed in real multi-omics data.

For each of the nine datasets, the resulting data maintained realistic intra- and inter-omics relationships while exhibiting different levels of cluster distinctness and noise. The generated data preserved key biological relationships, including correlations between CpG sites within the same CpG-island, the inverse relationship between upstream CpG methylation and gene expression (gene silencing), and the positive correlation between gene expression and downstream protein levels. This comprehensive synthetic data generation approach, utilizing different sample sizes and complexity levels, allowed us to systematically evaluate PEARL's performance and scalability under various controlled conditions with known ground-truth cluster assignments. The use of InterSIM ensured that our synthetic data maintained biological plausibility while providing the flexibility to test our method under different challenging scenarios.

#### Supplementary Note 3. Internal validation

##### Cross Validation and Performance Evaluation

To rigorously evaluate PEARL's performance and ensure robust statistical validation, we implemented a comprehensive cross-validation framework using StratifiedShuffleSplit. This approach maintains the original class distribution in both training and validation sets, which is particularly crucial for imbalanced datasets like BRCA and ROSMAP. The cross-validation process begins with an initial split of the dataset into training and test sets with an 8:2 ratio. The training set is then further divided into 30 different train-validation splits using StratifiedShuffleSplit, maintaining the same 8:2 ratio for each fold. This stratification ensures that the relative class frequencies are approximately preserved in each split, which is essential for maintaining the representativeness of the data.

The model configuration varies based on the dataset type. For binary classification tasks using the ROSMAP dataset, we employed an adjacency parameter of 2, hidden layer dimensions of [200, 200, 100], and SSGConv parameters of $K$ = 1 and $\alpha$ = 0.5. For multi-class problems, including both the BRCA dataset and synthetic datasets, we utilized an adjacency parameter of 10, hidden layer dimensions of [400, 400, 200], and SSGConv parameters of $K$ = 1 and $\alpha$ = 0.5. This sharing of configurations between BRCA and synthetic datasets reflects their similar multi-view nature.

The training process for each fold consists of two distinct phases. In the pretraining phase, the SSGConv layers are trained for 500 epochs with a learning rate of $1e-3$, focusing on optimizing the graph convolutional networks without the integration component. This is followed by the main training phase, where the complete model is trained for 2500 epochs with learning rates of $5e-4$ for the encoders and $1e-3$ for the classifier components. For each fold, we generate two types of adjacency matrices: training adjacency matrices using only the training data, and validation adjacency matrices that incorporate both training and validation data while maintaining proper separation between sets. This approach ensures that the graph structure accurately represents the relationships within each subset of the data while preventing information leakage between training and validation sets.

The performance evaluation metrics are tailored to the classification task at hand. For binary classification using the ROSMAP dataset, we compute accuracy, F1 score, and AUC score. For multi-class problems using BRCA and synthetic datasets, we calculate accuracy, weighted F1 score, and macro F1 score. These metrics are chosen to provide a comprehensive assessment of model performance across different aspects of classification quality. The results across all 30 folds are aggregated to compute mean performance metrics and standard deviations. Statistical significance is assessed through paired t-tests comparing PEARL's performance with competing methods. This comprehensive evaluation framework ensures the robustness of our results through multiple data partitions, maintains stability through stratified sampling, and guarantees reproducibility through fixed random seeds. By maintaining consistent evaluation procedures across all compared methods while adapting the model configuration to the specific characteristics of each dataset type, this framework provides a fair and thorough assessment of PEARL's performance. The use of 30 folds provides a reliable estimate of model performance variability, while the stratification in the splitting process ensures that the evaluation accurately reflects the model's performance across different class distributions.

To ensure optimal model performance, we implemented a comprehensive grid search methodology combined with cross-validation for both the ROSMAP and BRCA datasets. For each dataset, we explored different ranges of hyperparameters based on their specific characteristics and complexity. The complete parameter grids for ROSMAP and BRCA are detailed in **Supplementary Table 1** and **Supplementary Table 2**, respectively.

We implemented a rigorous 30-fold cross-validation procedure using StratifiedShuffleSplit to maintain class distribution consistency. For each fold, the data was initially divided into training-validation (70%) and test (30%) sets, with the training-validation portion further split into training (80%) and validation (20%) subsets. For each hyperparameter combination, the model was trained on the training set, with performance evaluated on the validation set. For ROSMAP, the AUROC served as the validation metric, while accuracy was used for BRCA. The best-performing parameter configuration on the validation set was then evaluated on the test set to obtain unbiased performance metrics. The results of the grid search are shown in **Supplementary Figure** 1. This comprehensive grid search strategy, combined with stratified cross-validation, ensured robust parameter selection while maintaining the integrity of the test set for final performance evaluation. The process was implemented using PyTorch, with all computations performed on GPU hardware when available for computational efficiency.

**Ablation Study**

To evaluate the impact of our key innovations, we conducted a comprehensive ablation study comparing four model variants. The complete PEARL model incorporates all three components, utilizing weighted Pearson correlation for similarity computation, SSGConv for graph convolution, and our adaptive integration method. PEARL-A maintains both the weighted Pearson correlation and SSGConv architecture but replaces our adaptive integration method with MOGONET's VCDN integration. PEARL-AP retains only the SSGConv architecture, replacing both the weighted Pearson correlation with cosine similarity and using MOGONET's VCDN integration. Finally, we included the original MOGONET as a baseline, which uses traditional graph convolution, cosine similarity, and VCDN integration. We evaluated these variants using the same cross-validation framework described previously on both ROSMAP and BRCA datasets.

On the ROSMAP dataset, we observed a clear hierarchical improvement with each innovation. While PEARL-A achieved slightly higher accuracy than the complete PEARL model, the complete PEARL still maintained superior performance compared to PEARL-AP and MOGONET. Importantly, the complete PEARL model demonstrated the highest performance in F1 score and AUROC, which are particularly critical metrics for binary classification tasks. This minor trade-off in accuracy was intentionally introduced to optimize the model's performance on multi-view datasets, as accuracy alone can be misleading in unbalanced datasets.

The BRCA dataset, however, revealed a more complex pattern. While PEARL still achieved the best performance across all metrics (Accuracy, F1 weighted, and F1 macro), the ordering of the other variants differed substantially. PEARL-AP performed second-best, followed by MOGONET, with PEARL-A showing the lowest performance among the variants. This unexpected ordering can be attributed to our development process, as the adaptive integration method was initially optimized on the ROSMAP dataset without considering the multi-class nature of the BRCA dataset. However, the consistent improvement of PEARL-AP over MOGONET across both datasets validates the effectiveness of SSGConv in handling both binary and multi-class scenarios. These findings ultimately led to the development of PEARL's final form, which successfully integrates all three innovations while maintaining robust performance across different classification tasks. The results are shown in **Supplementary Figure** 2.

The ablation study revealed crucial insights about our model components. The SSGConv architecture demonstrated consistent improvement in performance across both datasets, validating its effectiveness in capturing complex molecular interactions. The weighted Pearson correlation showed particular strength in the binary classification setting, while the adaptive integration method exhibited dataset-specific behavior, performing exceptionally well on binary classification tasks but requiring additional refinement for multi-class problems. Through these observations, we developed the final PEARL model to successfully address the limitations revealed in the ablation study, achieving superior performance across both binary and multi-class classification tasks. This comprehensive evaluation not only validates our design choices but also provides valuable insights into the interaction between different components of the model across varying classification scenarios. The results underscore the importance of careful component selection and adaptation based on the specific requirements of different classification tasks, particularly when handling multi-omics data with varying complexity and class structures.

#### Supplementary Tables

#### Supplementary Figures

| Parameter | Values |
|---|---|
| Hidden layer dimensions | [200, 200, 100], [250, 250, 125] |
| $K$ (SSGConv) | 1, 2, 3 |
| $\alpha$ (SSGConv) | 0.45, 0.5, 0.55 |
| Adjacency parameter | 2 |
| Dropout rate | 0.5 |
| Encoder learning rate | 1e-4, 5e-4 |
| Classifier learning rate | 1e-4, 5e-4 |
| Pretraining epochs | 500 |
| Training epochs | 2500 |

**Supplementary Table 1.** Hyperparameter Grid Configuration for ROSMAP dataset

| Parameter | Values |
|---|---|
| Hidden layer dimensions | [400, 400, 200], [500, 500, 250] |
| $K$ (SSGConv) | 1, 2, 3 |
| $\alpha$ (SSGConv) | 0.45, 0.5, 0.55 |
| Adjacency parameter | 10 |
| Dropout rate | 0.5 |
| Encoder learning rate | 1e-4, 5e-4 |
| Classifier learning rate | 1e-4, 5e-4 |
| Pretraining epochs | 500 |
| Training epochs | 2000 |

**Supplementary Table 2.** Hyperparameter Grid Configuration for BRCA dataset

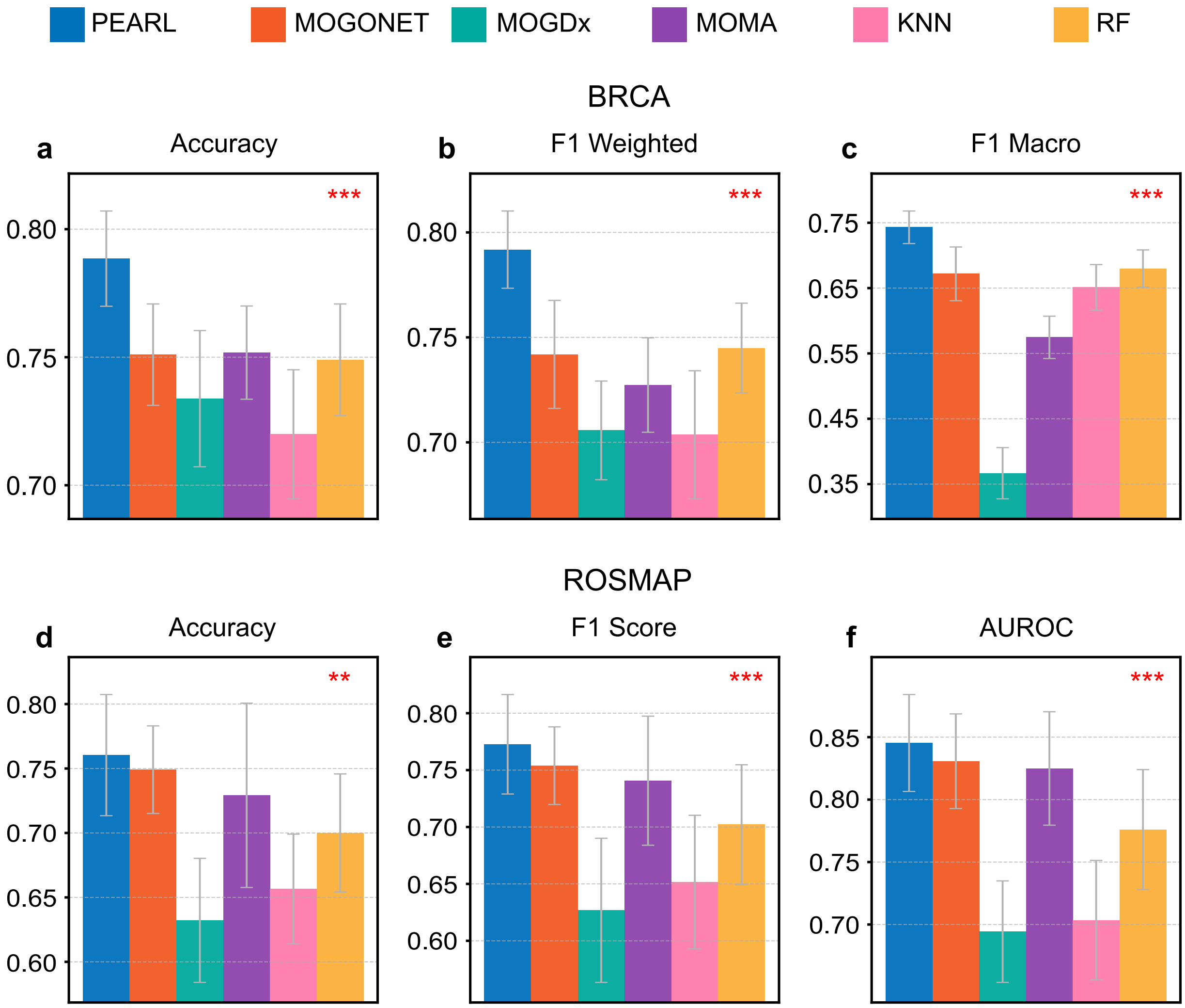


**Supplementary Figure 1. Grid search on real datasets.** We compared the performance of PEARL with four other methods, MOGONET, MOGDx, MOMA, KNN, and RF, on the BRCA dataset and the ROSMAP dataset. We used Accuracy, F1 Weighted, and F1 Macro as metrics on the BRCA multi-view dataset. We used Accuracy, F1 Score, and AUROC as metrics on the ROSMAP binary dataset. We conducted paired t-tests comparing PEARL with the second-best method in each case. The resulting $P$-values are indicated with asterisks in the plots, where three asterisks denote a $P$-value less than 0.01, and two asterisks denote a $P$-value less than 0.05.

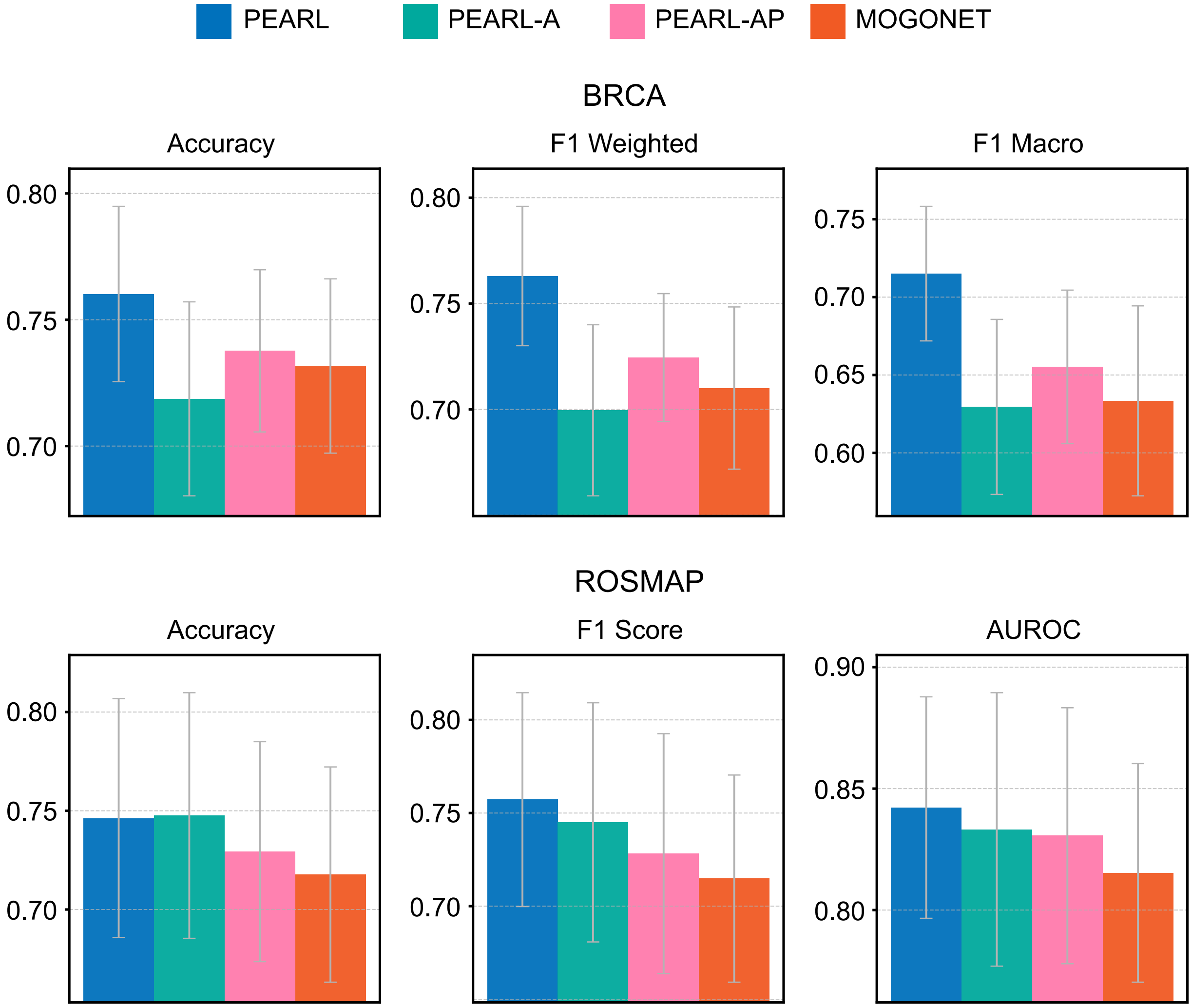


**Supplementary Figure 2. Ablation study results comparing different model variants.** Performance comparison between PEARL and its variants on the ROSMAP (binary classification) and BRCA (multi-class classification) datasets. PEARL represents our complete model with all three key innovations: the use of SSGConv, weighted Pearson correlation, and adaptive integration method. PEARL-A retains SSGConv and weighted Pearson correlation but uses MOGONET's VCDN integration. PEARL-AP keeps only SSGConv, using MOGONET's cosine similarity and VCDN integration. MOGONET represents the baseline model with traditional graph convolution, cosine similarity, and VCDN integration. We used Accuracy, F1 Weighted, and F1 Macro as metrics on the BRCA multi-view dataset. We used Accuracy, F1 Score, and AUROC as metrics on the ROSMAP binary dataset.

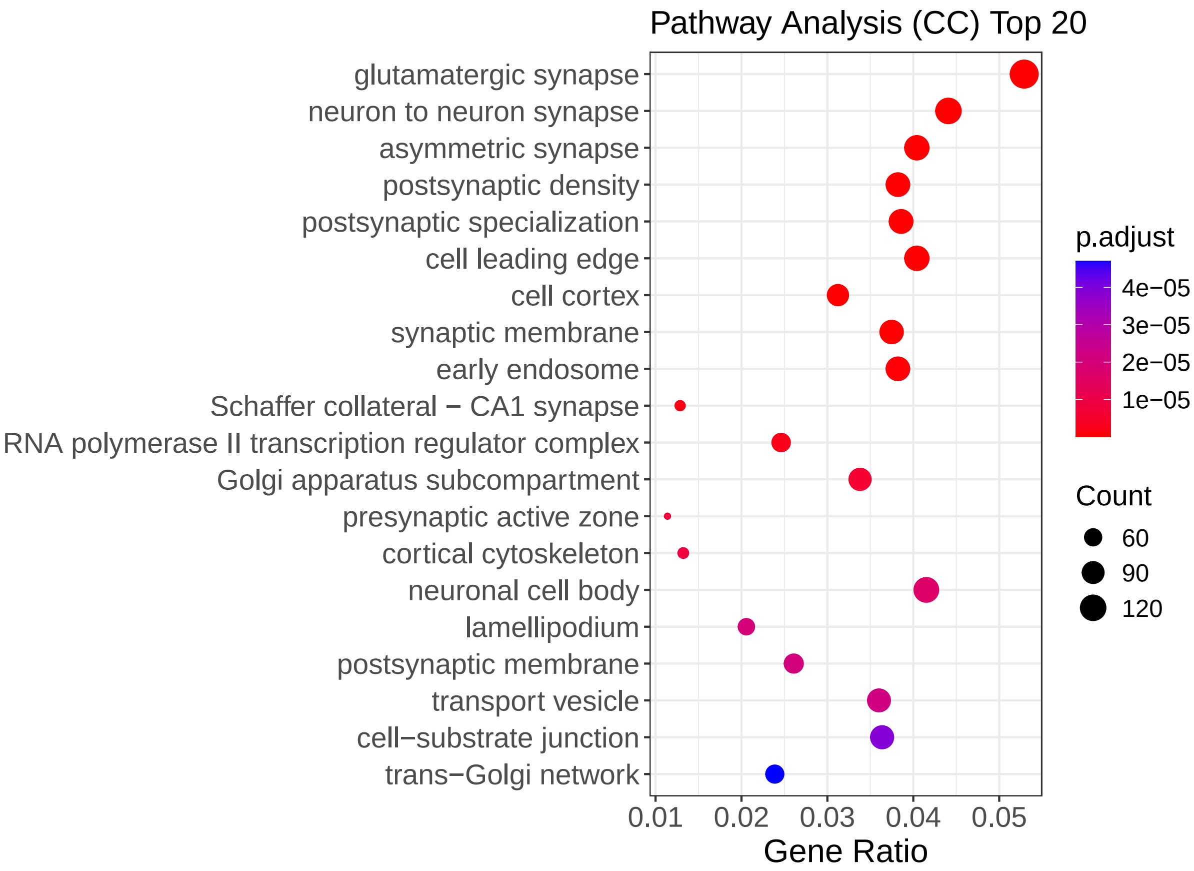


**Supplementary Figure 3. Top 20 Enriched Cellular Components from Pathway Analysis.** Top 20 enriched pathways ordered by adjusted $P$-value. Each dot represents a cellular component, with the x-axis showing the gene ratio (the ratio of genes associated with a given term to the total number of genes analyzed), and the y-axis listing the GO term descriptions. Color corresponds to the adjusted $P$-value, with more intense red indicating higher significance. Dot size reflects the gene count.

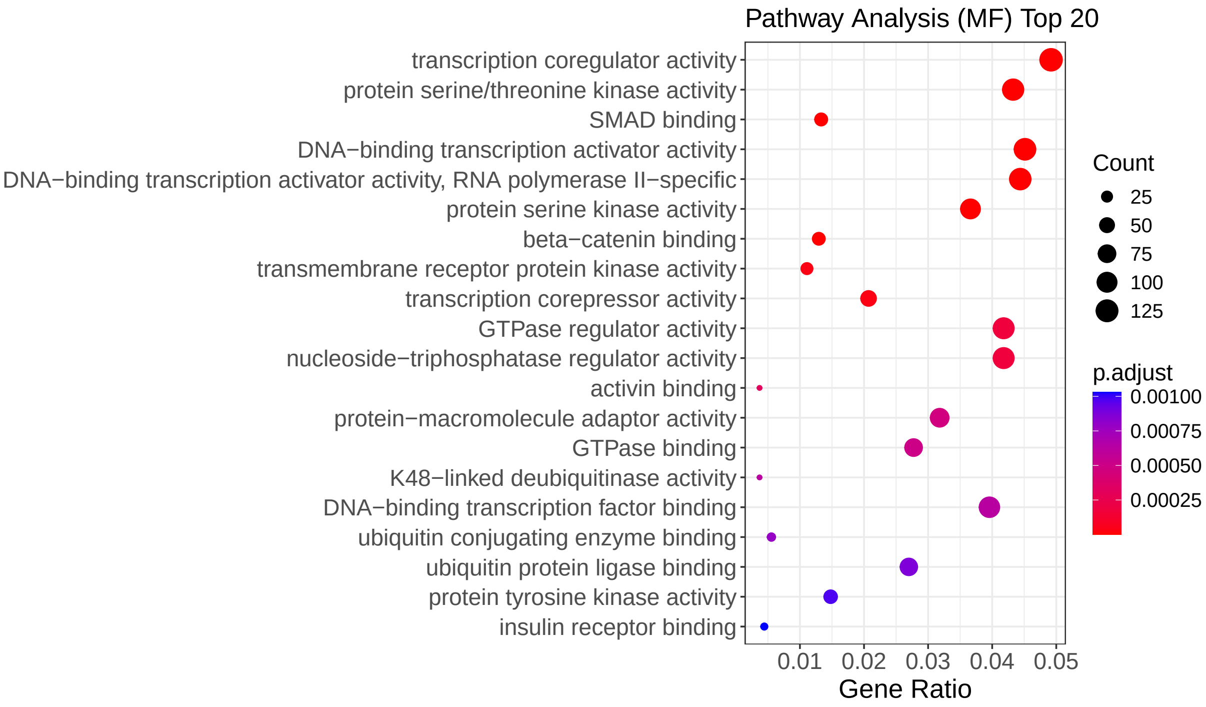


**Supplementary Figure 4. Top 20 Enriched Molecular Function from Pathway Analysis.** Top 20 enriched pathways ordered by adjusted $P$-value. Each dot represents a molecular function, with the x-axis showing the gene ratio (the ratio of genes associated with a given term to the total number of genes analyzed), and the y-axis listing the GO term descriptions. Color corresponds to the adjusted $P$-value, with more intense red indicating higher significance. Dot size reflects the gene count.

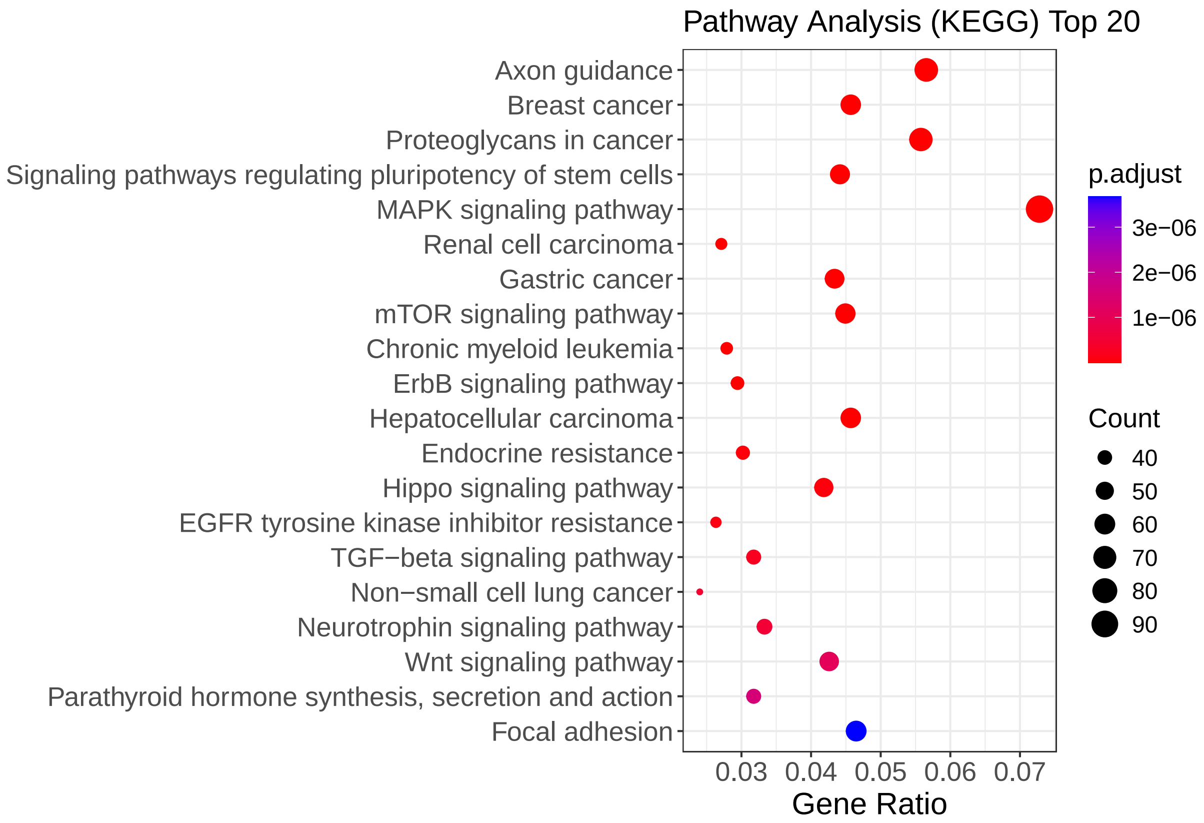


**Supplementary Figure 5. Top 20 Enriched KEGG pathways.** Top 20 enriched pathways ordered by adjusted $P$-value. Each dot represents a KEGG pathway, with the x-axis showing the gene ratio (the ratio of genes associated with a given term to the total number of genes analyzed), and the y-axis listing the KEGG term descriptions. Color corresponds to the adjusted $P$-value, with more intense red indicating higher significance. Dot size reflects the gene count.
